# Gut microbiome response to a modern Paleolithic diet in a Western lifestyle context

**DOI:** 10.1101/494187

**Authors:** Monica Barone, Silvia Turroni, Simone Rampelli, Matteo Soverini, Federica D’Amico, Elena Biagi, Patrizia Brigidi, Emidio Troiani, Marco Candela

**Affiliations:** Unit of Microbial Ecology of Health, Department of Pharmacy and Biotechnology, University of Bologna, Italy; Primary Care Unit and Territorial Health, Social Security Institute, Republic of San Marino.

## Abstract

The progressive reduction of gut microbiome (GM) biodiversity along human evolutionary history has been found to be particularly exacerbated in Western urban compared to traditional rural populations, and supposed to contribute to the increasing incidence of chronic non-communicable diseases. Together with sanitation, antibiotics and C-section, the Western diets, low in microbiota-accessible carbohydrates (MACs) while rich in industrialized and processed foods, are considered one of the leading causes of this shrinkage. However, significant questions remain unanswered, especially whether high-MAC low-processed diets may be sufficient to recover GM diversity in Western urban populations. Here, we profiled the GM structure of urban Italian subjects adhering to the modern Paleolithic diet (MPD), a dietary pattern featured by high consumption of MACs and low-to-zero intake of refined sugars and processed foods, and compared data with other Italian individuals following a Mediterranean Diet (MD), as well as worldwide traditional hunter-gatherer populations from previous publications. Notwithstanding a strong geography effect on the GM structure, our results show an unexpectedly high degree of GM biodiversity in MPD subjects, which well approximates that of traditional populations. Increasing the consumption of MACs at the expense of refined sugars, and minimizing the intake of processed foods, both hallmarks of the MPD, could be the key to rewild the Western microbiota, counteracting the loss of GM diversity and thus restoring evolutionarily important functionality to our gut for improved human health.

## Introduction

In order to understand the specificities of the human microbiome assembly, extensive meta-analyses of human and non-human primate microbiomes have been recently carried out [1,2]. This comparative approach has highlighted the reduction of individual biodiversity as one of the distinctive features of the human gut microbiome (GM) [1]. Interestingly, this hallmark has been found to be exacerbated in Western urban populations, which show even more marked compression of personal diversity than traditional and rural counterparts [3–6]. Consistent with the disappearing microbiota hypothesis [7], the dramatic shrinkage of individual GM diversity in Western urban populations depicts a maladaptive microbiome state that has been supposed to contribute to the rising incidence of chronic non-communicable diseases, such as obesity, diabetes, asthma and inflammatory bowel disease [8–11]. Consequently, in recent years, a large body of research has been devoted to understanding the mechanisms leading to the diversity loss in the Western urban GM. It is in this scenario that the multiple-hit hypothesis has been advanced [8]. According to this theory, the progressive reduction of human GM diversity has occurred at multiple stages along the recent transition to modern urban societies, and several aspects typical of the urbanization process - such as sanitation, antibiotics, C-section and Western diet - have been pointed out as contributing factors. In particular, the reduction in quantity and diversity of Microbiota-Accessible Carbohydrates (MACs) in the diet has been considered one of the leading causes of the disappearing GM in Western urban populations [8]. Recently defined, MACs include fermentable fibers that - indigestible by the host - become available as an energy source for a specific GM fraction enriched in Carbohydrate Active Enzymes (CAZymes) [8,12]. Moreover, food additives, emulsifiers and xenobiotics – ubiquitous in industrially processed foods – have recently been shown as important additional drivers of GM diversity shrinkage [13].

All currently available studies exploring the disappearing GM are based on the comparison between Western urban and traditional rural populations [3–6,14–16]. Consistently, the observed GM differences are likely to be the result of the combined action of several covariates in addition to the diet – i.e. ethnicity, geographical origin, climate, subsistence, medication, hygiene and life sharing – and do not allow to weight the importance of the single determinants. Indeed, to the best of our knowledge, no study has been specifically designed to dissect the role of MACs deprivation and xenobiotics exposure as driving factors forcing the compression of GM diversity in Western urban populations.

In the last few years, the Modern Paleolithic Diet (MPD), with high intake of vegetables, fruit, nuts, eggs, fish and lean meat, while excluding grains, dairy products, salt and refined sugar, has attracted substantial public attention in the Western world because of its potential multiple health benefits [17–20]. Being enriched in MACs and completely excluding industrially processed food, the MPD pattern represents a model of Western urban diet ideal for disentangling the impact of MACs and food xenobiotics on the human GM.

In the present work, we profiled the GM structure of 15 Italian subjects following the MPD and compared it with that of Italian individuals largely adhering to the Mediterranean Diet (MD) from our previous works [5,21]. Notwithstanding the small sample size, our GM exploratory study gave us the unique opportunity to assess to what extent the consumption of high amounts of MACs along with the exclusion of industrially processed food, may counteract the GM diversity reduction as observed in Western urban populations. Indeed, the comparison between MPD and Western diets in subjects living in the same country allows excluding the impact of confounding drivers of GM variation, such as geography, ethnicity, medication, hygiene and subsistence [14,15,21]. In order to extend the GM comparison at the meta-population level, we expanded our meta-analysis by including publically available microbiome data from traditional hunting and gathering populations showing high GM diversity, such as the Hadza from Tanzania, from our previous publication [5], the Matses from Peru [6], and the Inuit from the Canadian Arctic [22].

According to our data, increased consumption of unprocessed foods and dietary MACs, as observed in MPD individuals, is sufficient, even in a Western context, to recover the levels of GM diversity typically found in traditional rural populations. Although the mechanisms underlying the human-microbiome interactions are still far from being fully understood, the possibility of rewilding the modern microbiota through the diet could be the key to restore evolutionarily important functionality to the gut, ultimately improving our health.

## Results

### Diet and gut microbiome structure in Italian adults following the modern Paleolithic diet

Fifteen healthy individuals who have been following the MPD for at least one year were recruited from different urban areas across Italy (Lombardia, Piemonte, Emilia-Romagna, Toscana, Umbria, Lazio, Campania, Molise, Puglia and Calabria regions). Specifically, 3 female and 12 male adults, with an average age of 39.2 years (range, 26 – 57) and average Body Mass Index (BMI) of 22.1 kg·m^-2^ (range, 19.4 – 25.7), were enrolled in our cohort (S1 Table).

The MPD adopted by the 15 subjects is mainly based on the consumption of unprocessed foods, with high intake of vegetables, fruit, nuts, eggs, fish and lean meat, while excluding grains, dairy products, salt and refined sugar. The daily total calorie intake, as well as that of macro- and micronutrients, assessed through 7-day weighted food intake records (7D-WRs), are reported in S2 Table. The average daily energy intake of the enrolled cohort is 1,843.45 kcal (range, 1,563 – 2,186 kcal). The percentage of macronutrients is distributed as follows: fat, 51.02%; protein, 30.14%; carbohydrate, 18.84% (Fig 1A). With regard to lipids, 51.65% of total calories are from monounsaturated fatty acids (MUFAs), 30.93% from saturated fatty acids (SFAs) and 17.42% from polyunsaturated fatty acids (PUFAs) (Fig 1B). The average daily fiber intake is 14.64 g/1000 kcal. The GM structure of MPD Italian adults was profiled through 16S rRNA gene sequencing of fecal DNA. A total of 864,439 high-quality reads (mean ± sd, 57.6 ± 19.7; range, 25,142 – 95,924) were generated and clustered in 7,483 OTUs. The phyla Firmicutes (relative abundance, mean ± sem, 65.1 ± 2.1 %) and Bacteroidetes (24.6 ± 2.2 %) dominate the gut microbial ecosystem, with Proteobacteria (4.4 ± 1.6 %), Actinobacteria (3.4 ± 0.8%) and Verrucomicrobia (1.2 ± 0.5 %) as minor components. At family level, *Ruminococcaceae* (26.7 ± 1.7 %), *Lachnospiraceae* (18.7 ± 1.4 %), *Bacteroidaceae* (13.7 ± 1.8 %) and *Prevotellaceae* (7.4 ± 2.4 %) are the dominant GM constituents. The most abundant (≥ 5%) bacterial genera are *Bacteroides, Prevotella*, and *Faecalibacterium*, while *Coprococcus, Ruminococcus, Blautia, Lachnospira, Phascolarctobacterium, Streptococcus, Roseburia, Akkermansia, Oscillospira* and *[Eubacterium]* represent minor components of the microbial ecosystem (range 4.4 ± 0.7 % - 1.0 ± 0.4 %) (Fig 2).

**Fig 1.**
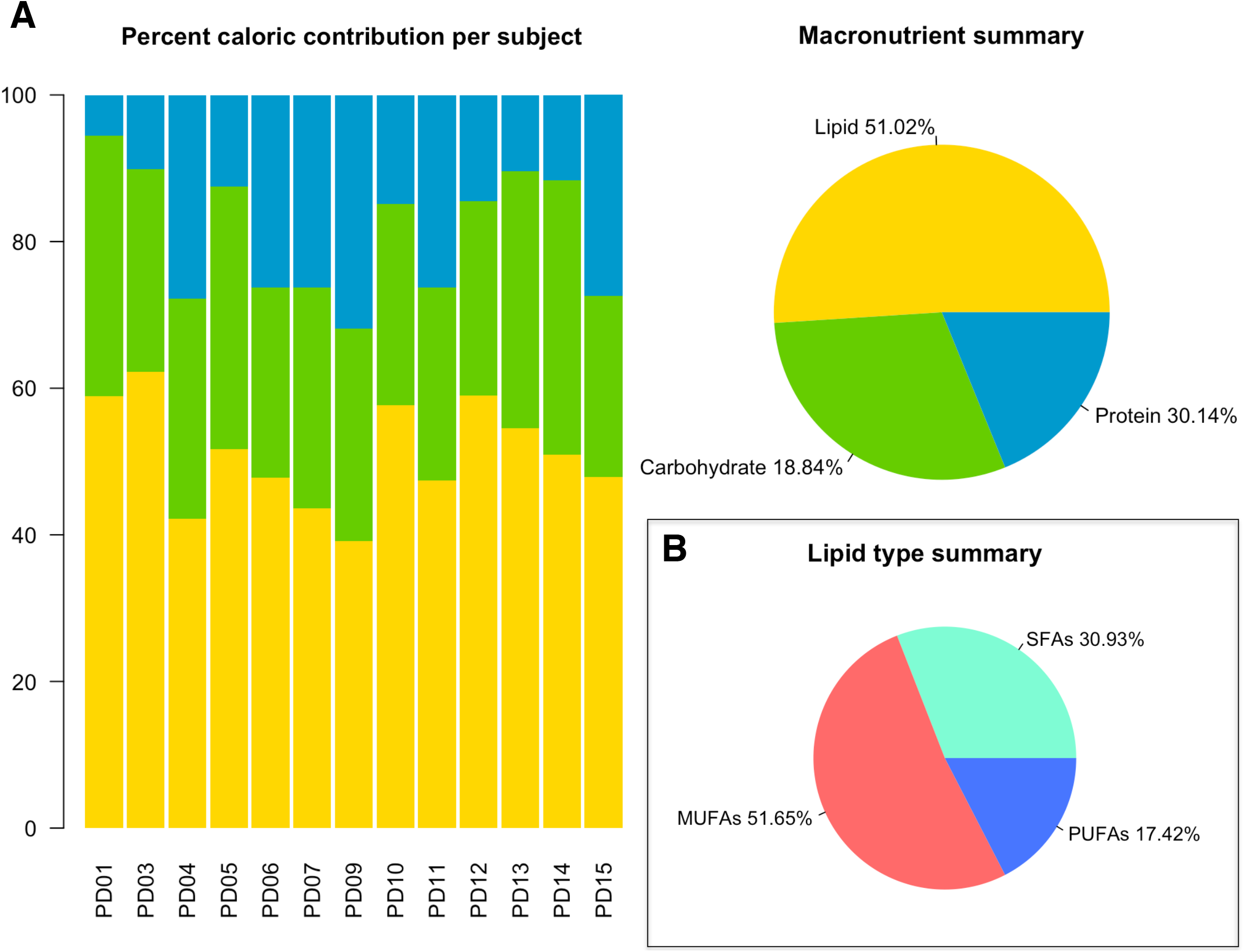
Macronutrient composition of the modern Paleolithic diet. (A) Bar plots of the percent caloric contribution of fat, protein and carbohydrate per subject, based upon weighted food intake records over 7 days. The pie chart shows the summary of the average macronutrient intake for the entire cohort. (B) Pie chart of the lipid type summary. PUFAs: polyunsaturated fatty acids; MUFAs: monounsaturated fatty acids; SFAs: saturated fatty acids.

**Fig 2.**
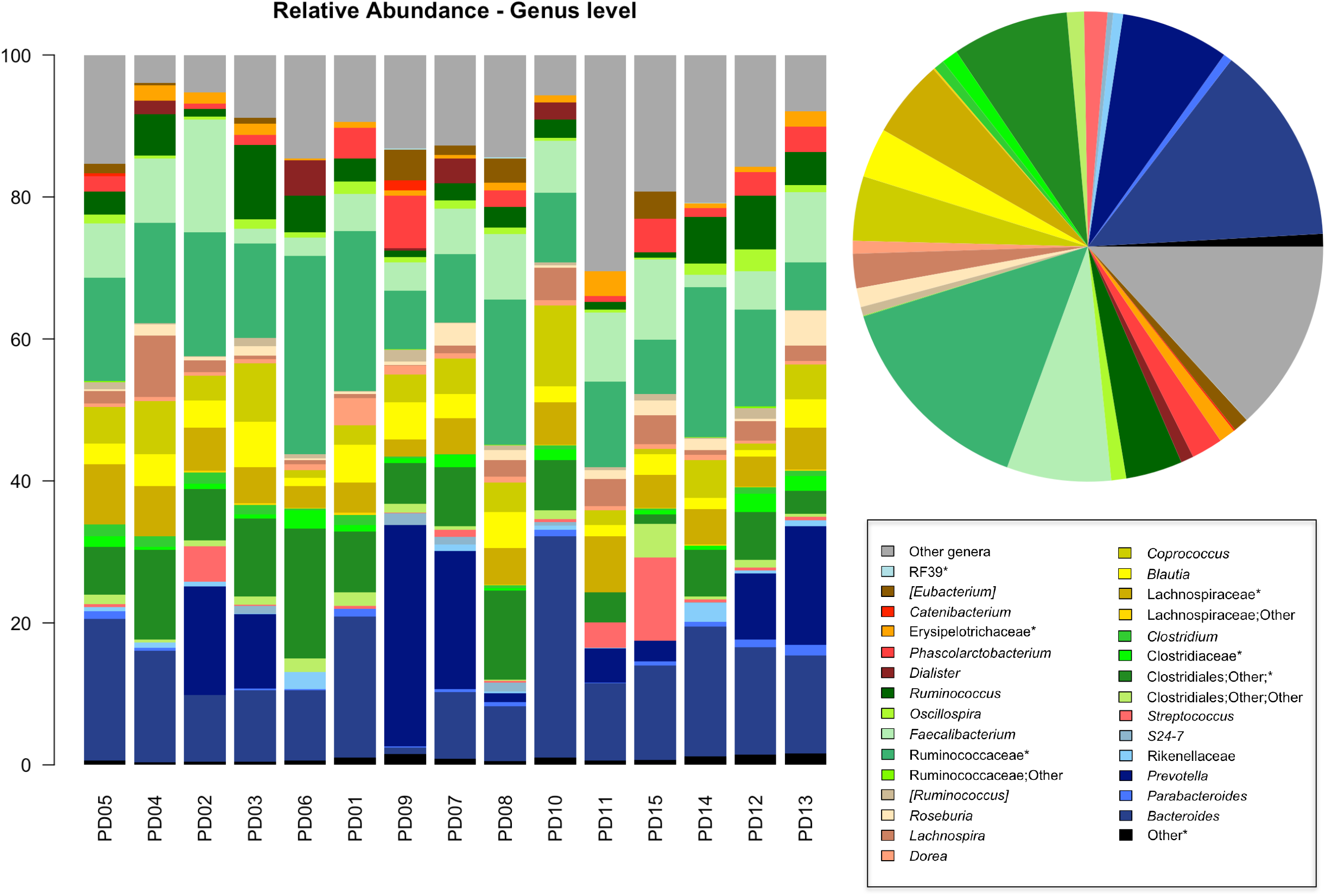
Phylogenetic structure of the gut microbiome of Italian adults adhering to the modern Paleolithic diet. Bar plots of the genus-level composition of the gut microbiome of the enrolled subjects. The pie chart shows the average relative abundance of bacterial genera. Only bacterial genera with relative abundance > 0.5 % are shown. * = Unclassified.

### Gut microbiome diversity in MPD Italian adults and comparison with other Western urban populations and traditional communities

In order to investigate whether the adherence to the MPD is sufficient to promote a more diverse GM ecosystem - even in a Western urban context - we compared the GM diversity between the 15 MPD subjects and 143 urban Italians with different level of adherence to the MD, whose GM composition was described in De Filippis *et al.* [21] and Schnorr *et al.* [5]. Moreover, to extend the meta-analysis to a global level, the GM structural profiles of the following traditional hunter-gatherer populations were included: 27 Hadza from Tanzania [5], 25 Matses from Peru [6], and 21 Inuit from Canada [22]. According to our findings, significant differences in the GM biodiversity among the study groups were detected (Simpson index, P-value = 2.6 × 10^-15; Shannon index, P-value = 2.2 × 10^-16; Kruskal-Wallis test) (Fig 3). Interestingly, the GM diversity observed for MPD subjects far exceeds that of urban Italians adhering to the MD (Simpson index, P-value = 2.5 x 10^-7^; Shannon index, P-value = 6.1 x 10^-9^; Wilcoxon test), and is even greater than that observed in Matses (P-value = 0.0082; 0.0039) and Inuit (P-value = 0.00075; 0.0027). On the other hand, no significant difference was found between MPD individuals and the Hadza (P-value = 0.39; P-value = 0.26).

**Fig 3.**
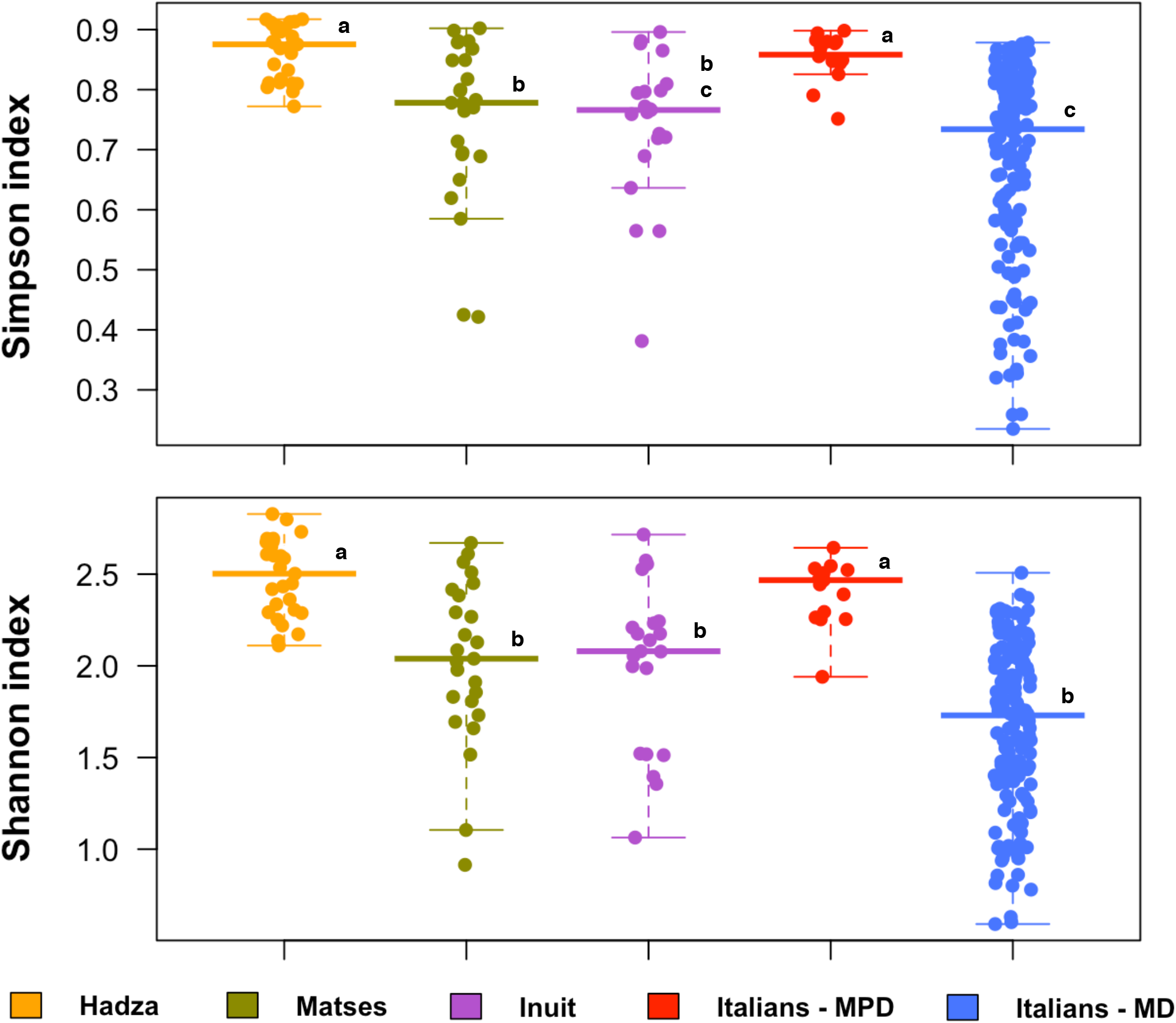
The gut microbiome of Italian subjects following the modern Paleolithic diet shows intermediate biodiversity between Western urban and traditional populations. Box and scatter plots showing the alpha diversity values, measured with Simpson and Shannon indices, for each study population (i.e. urban Italians adhering to the modern Paleolithic diet from the present study, urban Italians adhering to the Mediterranean diet [21], Hadza from Tanzania [5], Matses from Peru [6], and Inuit from Canadian Arctic [22]). Different letters in the box plots indicate significant differences (P-value < 0.05, Wilcoxon test). MPD = Modern Paleolithic Diet; MD = Mediterranean Diet.

The PCoA based on Bray-Curtis distances was next used to assess overall genus-level compositional differences in the GM structure between study groups. Our data show clear separation of GM profiles by provenance and, within the Italian cohort, by dietary pattern (P-value < 1 × 10^-5^, permutation test with pseudo-F ratios) (Fig 4A). Interestingly, MPD subjects show a low level of interpersonal GM variation (Bray-Curtis distances, mean ± SD, 0.42 ± 0.095), approximating that observed for the Hadza (0.36 ± 0.092) (Fig 4B). Finally, to identify the bacterial drivers with a statistically significant contribution (permutation correlation test, P-value < 0.001) to the sample ordination, we superimposed the genus relative abundance on the PCoA plot (S1 Fig). According to our data, the microorganisms characterizing the Italian cohort are *Bacteroides, Collinsella, Coprococcus* and *Blautia.* The genera *Clostridium, Prevotella, [Prevotella], Catenibacterium* and *Oscillospira* were found to be associated with Hadza and Matses, while *Sutterella* and *Parabacteroides* with Inuit.

**Fig 4.**
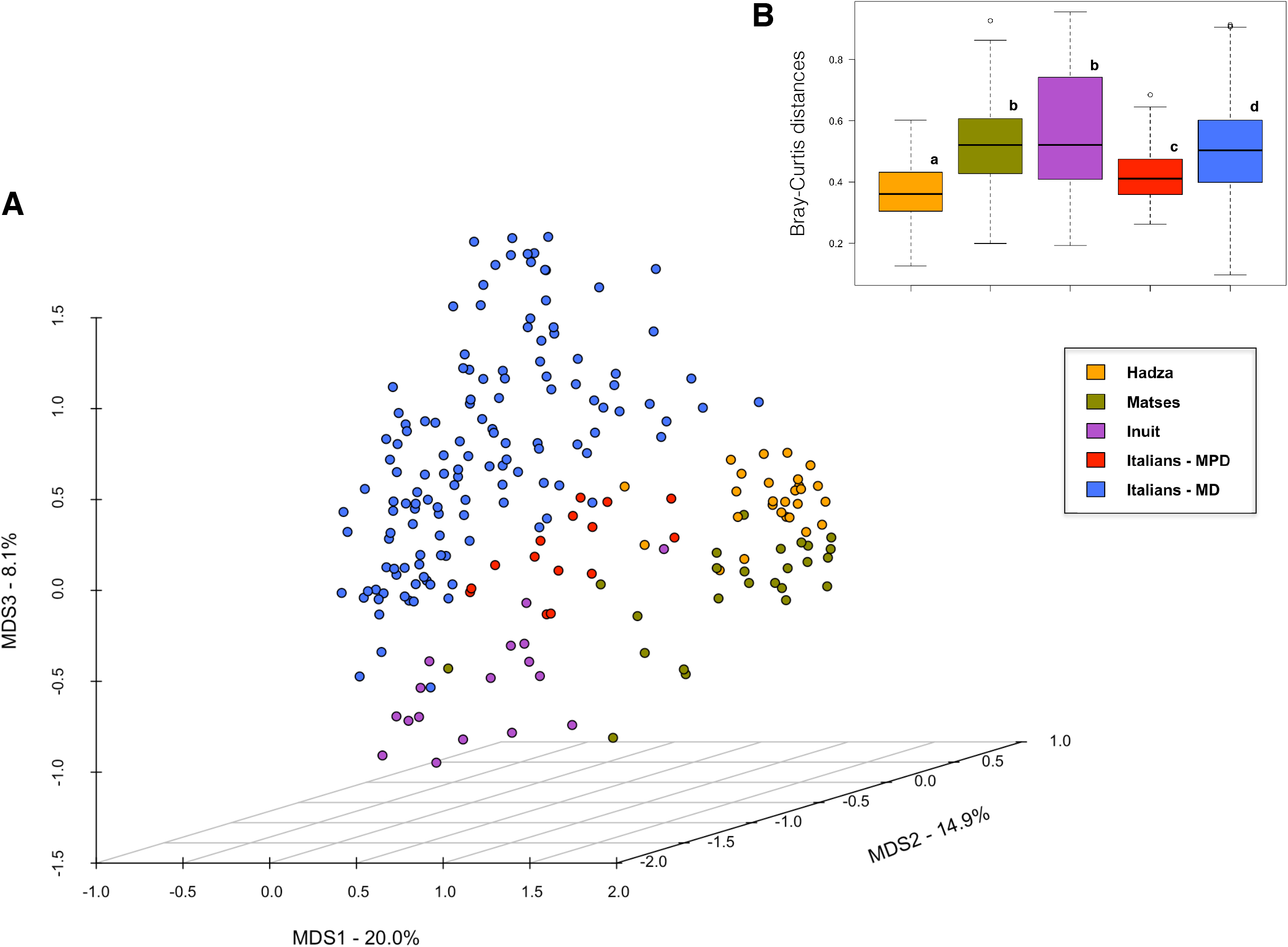
Beta diversity of the fecal microbiome of Italian subjects following the modern Paleolithic diet compared with other Western urban populations and traditional communities. (A) The PCoA plot shows the Bray-Curtis distances between the genus-level microbiota profiles of urban Italians adhering to the modern Paleolithic diet from the present study, urban Italians adhering to the Mediterranean diet [21], Hadza from Tanzania [5], Matses from Peru [6], and Inuit from Canadian Arctic [22]. A significant segregation among study populations was found (P-value < 1 × 10^-5^; permutation test with pseudo-F ratios). (B) Boxplots represent the interpersonal variation, in terms of Bray-Curtis distances between genus-level microbiota profiles. Different letters in the boxplots indicate significant differences (P-value < 0.05, Wilcoxon test). MPD = Modern Paleolithic Diet; MD = Mediterranean Diet.

## Discussion

Herein we compared the GM compositional structure and diversity of 15 urban Italian adults adhering to the MPD with previously published data from 143 urban Italian adults largely adhering to the MD [5,21] and 73 traditional hunter-gatherers, including 27 Hadza from Tanzania [5], 25 Matses from Peru [6], and 21 Inuit from Canada [22]. According to our findings, the study groups segregate by geographical origin, with a further separation within the Italian cohort reflecting the diet pattern (MPD vs MD). This provenance-dependent effect on the human GM structure probably involves the concomitant action of several covariates, which concur in shaping the GM structure, such as geography, ethnicity, lifestyle and dietary habits. The Italian origin of the GM seems to be defined by a higher abundance of *Bacteroides, Collinsella, Coprococcus* and *Blautia*, bacterial genera commonly found within Western microbiomes [3–6]. According to the literature, the separation due to geography seems to be less evident among the traditional populations, with Matses and Hadza sharing a high abundance of *Prevotella* [5,6].

These data confirm recent findings that demonstrate the predominance of host location and ethnicity, with respect to diet, as determinants of human GM variation [14,15]. However, despite the overall Western-like configuration, the MPD-associated GM structure stands out from that of Italians adhering to the MD for a much higher degree of biodiversity, which well approximates that observed in traditional hunter-gatherers. Since the Italian subjects of our cohort share the provenance and all that it entails, including the lifestyle, it can be hypothesized that the MPD-associated bloom in GM diversity is accounted for by the peculiarities of the MPD compared to the MD. Though the two diets are similar in many respects – i.e. high intake of fruit, vegetables, fish and nuts, as well as low glycemic load – the MPD is in fact distinguished by: (i) greater consumption of MACs, mainly from wild plant foods; (ii) total exclusion of industrially processed products; (iii) higher intake of unsaturated fatty acids, especially MUFAs, from olive oil, nuts and meat; (iv) no consumption of foods containing refined sugars [17–20]. It is, therefore, tempting to speculate that these MPD distinctive features may be sufficient to support the consolidation of a highly diversified GM layout, thus counteracting the loss of GM biodiversity, typically observed in Western urban populations as compared to traditional communities [3–6].

In conclusion, our findings shed some light on the possibility of recovering GM diversity in Western urban populations through diet. Increasing the consumption of MACs at the expense of refined sugars, and minimizing the intake of processed foods, both hallmarks of the MPD, could indeed act synergistically in facilitating the regain of an eubiotic level of GM diversity. Moreover, the high intake of MUFAs, as found in the MPD, suggests that these fatty acids could play a role in supporting the increase in GM diversity, which is worthy of being further explored in larger cohorts.

## Materials and methods

### Subjects and sample collection

Fifteen healthy individuals following a MPD for at least one year were recruited from different urban areas across Italy (Lombardia, Piemonte, Emilia-Romagna, Toscana, Umbria, Lazio, Campania, Molise, Puglia and Calabria regions). Anthropometric data and stool samples were collected from each participant who had not taken antibiotics in the previous 3 months. Fecal samples were immediately frozen at -20°C and then delivered to the laboratory of the Microbial Ecology of Health Unit (Dept. Pharmacy and Biotechnology, University of Bologna, Bologna, Italy) where they were stored at -80°C until processing. Each subject was asked to fill in a 7-day weighted food intake record (7D-WR), with the total food and beverage consumption, as previously described [23]. Daily total calorie intake as well as that of macro- and micro-nutrients were assessed through the MètaDieta^®^ software version 3.7 (METEDA). Written informed consent was obtained from all volunteers. All work was approved by the Ethics Committee of the Sant’Orsola-Malpighi Hospital, University of Bologna (ref. number, 118/2015/U/Tess).

### Microbial DNA extraction

Total bacterial DNA was extracted from each stool sample using the DNeasy Blood and Tissue kit (QIAGEN) with the modifications previously described by Biagi *et al.* [24]. In brief, 250 mg of fecal samples were suspended in 1 ml of lysis buffer (500 mM NaCl, 50 mM Tris-HCl pH 8, 50 mM EDTA, 4% (w/v) SDS), added with four 3-mm glass beads and 0.5 g of 0.1-mm zirconia beads (BioSpec Products) and homogenized using a FastPrep instrument (MP Biomedicals) with three bead-beating steps at 5.5 movements/sec for 1 min, and 5-min incubation in ice between treatments. After incubation at 95°C for 15 min, stool particles were pelleted by centrifugation at 14,000 rpm for 5 min. Nucleic acids were precipitated by adding 260 μl of 10 M ammonium acetate and one volume of isopropanol. The pellets were then washed with 70% ethanol and suspended in TE buffer. RNA was removed by treatment with 2 μl of DNase-free RNase (10 mg/ml) at 37°C for 15 min. Protein removal and column-based DNA purification were performed following the manufacturer’s instructions (QIAGEN). DNA was quantified with the NanoDrop ND-1000 spectrophotometer (NanoDrop Technologies).

### 16S rRNA gene sequencing

For each sample, the V3-V4 region of the 16S rRNA gene was amplified using the S-D-Bact-0341-b-S-17/S-D-Bact-0785-a-A-21 primers [25] with Illumina overhang adapter sequences. PCR reactions were performed in a final volume of 25 μl, containing 12.5 ng of genomic DNA, 200 nM of each primer, and 2X KAPA HiFi HotStart ReadyMix (Kapa Biosystems, Roche), in a Thermal Cycler T (Biometra GmbH) with the following gradient: 3 min at 95°C for the initial denaturation, 25 cycles of denaturation at 95°C for 30 sec, annealing at 55°C for 30 sec and extension at 72°C for 30 sec, and a final extension step at 72°C for 5 min. PCR products of around 460 bp were purified using a magnetic bead-based system (Agencourt AMPure XP; Beckman Coulter) and sequenced on Illumina MiSeq platform with the 2 × 250 bp paired-end protocol, according to the manufacturer’s guidelines (Illumina). Briefly, each indexed library was prepared by limited-cycle PCR using Nextera technology, and further purified as described above. The libraries were subsequently pooled at equimolar concentrations, denatured with 0.2 N NaOH, and diluted to 6 pM before loading onto the MiSeq flow cell. Sequencing reads were deposited in MG-RAST (ID: …).

### Bioinformatics and statistics

Raw sequences were processed using a pipeline that combines PANDAseq [26] and QIIME [27]. The UCLUST software [28] was used to bin high-quality reads into operational taxonomic units (OTUs) at 0.97 similarity threshold through an open-reference strategy. Taxonomy was assigned through the RDP classifier, using the Greengenes database as a reference (release May 2013). Chimera filtering was performed by using ChimeraSlayer [29]. All singleton OTUs were discarded. 16S rRNA gene sequencing data of our cohort were compared with publicly available data from the following previous studies: De Filippis *et al.* [21] (127 Italians; NCBI Sequence Read Archive (SRA) accession number: SRP042234), Schnorr *et al.* [5] (16 Italians and 27 Hadza hunter-gatherers from Tanzania; MG-RAST ID: 7058), Obregon-Tito *et al.* [6] (25 Matses hunter-gatherers from Peru; NCBI SRA: PRJNA268964), and Girard *et al.* [22] (21 Inuit from the Canadian Arctic; Qiita ID: 10439). Alpha diversity was assessed using the Shannon and Simpson indices. Beta diversity was evaluated using the Bray-Curtis dissimilarity measure. All statistical analysis was performed in R 3.3.2, using R Studio 1.0.44 and the libraries vegan, made4 and stats. The significance of data separation in the Principal Coordinates Analysis (PCoA) of Bray-Curtis distances was tested using a permutation test with pseudo-F ratios (function Adonis of vegan). Superimposition of bacterial genera on the PCoA plot was performed using the envfit function of vegan. Wilcoxon test was used to assess significant differences between groups (for intra- and interindividual diversity), while Kruskal–Wallis test was used for multiple comparisons. P-values were corrected for false discovery rate (FDR, Benjamini-Hochberg) and P-values ≤ 0.05 were considered statistically significant.

## Supporting information

## Acknowledgements

We would like to thank, and Jessica Baldini and Sara Quercia from Wellmicro S.r.l. (Bologna, Italy) for the skillful technical assistance in processing 7D-WRs and sequencing fecal samples, as well as the Paleo Meeting Organizing Committee (Italy).

## Supporting information

**S1 Fig. Superimposition of the genus relative abundance on the PCoA plot.** Arrows represent the direction of significant correlations (permutation correlation test, P-value < 0.001). MPD = Modern Paleolithic Diet; MD = Mediterranean Diet.

**S1 Table. Anthropometric data of the enrolled cohort.**

**S2 Table. MPD macro- and micro-nutrients summary based on MétaDieta output.**

## References

1. Moeller AH, Li Y, Mpoudi Ngole E, Ahuka-Mundeke S, Lonsdorf EV, Pusey AE, et al. Rapid changes in the gut microbiome during human evolution. Proc Natl Acad Sci U S A. 2014;111(46):16431–5.

2. Davenport ER, Sanders JG, Song SJ, Amato KR, Clark AG, Knight R. The human microbiome in evolution. BMC Biol. 2017;15(1):127.

3. De Filippo C, Cavalieri D, Di Paola M, Ramazzotti M, Poullet JB, Massart S, et al. Impact of diet in shaping gut microbiota revealed by a comparative study in children from Europe and rural Africa. Proc Natl Acad Sci U S A. 2010;107(33):14691–6.

4. Yatsunenko T, Rey FE, Manary MJ, Trehan I, Dominguez-Bello MG, Contreras M, et al. Human gut microbiome viewed across age and geography. Nature. 2012;486(7402):222–7.

5. Schnorr SL, Candela M, Rampelli S, Centanni M, Consolandi C, Basaglia G, et al. Gut microbiome of the Hadza hunter-gatherers. Nat Commun. 2014;5:3654.

6. Obregon-Tito AJ, Tito RY, Metcalf J, Sankaranarayanan K, Clemente JC, Ursell LK, et al. Subsistence strategies in traditional societies distinguish gut microbiomes. Nat Commun. 2015;6:6505.

7. Blaser MJ. The theory of disappearing microbiota and the epidemics of chronic diseases. Nat Rev Immunol. 2017;17(8):461–3.

8. Sonnenburg ED, Sonnenburg JL. Starving our microbial self: the deleterious consequences of a diet deficient in microbiota-accessible carbohydrates. Cell Metab. 2014;20(5):779–86.

9. Mosca A, Leclerc M, Hugot JP. Gut Microbiota Diversity and Human Diseases: Should We Reintroduce Key Predators in Our Ecosystem? Front Microbiol. 2016;7:455.

10. Zuo T, Kamm MA, Colombel JF, Ng SC. Urbanization and the gut microbiota in health and inflammatory bowel disease. Nat Rev Gastroenterol Hepatol. 2018;15(7):440–52.

11. Cani PD, Jordan BF. Gut microbiota-mediated inflammation in obesity: a link with gastrointestinal cancer. Nat Rev Gastroenterol Hepatol. 2018. [Epub ahead of print].

12. El Kaoutari A, Armougom F, Gordon JI, Raoult D, Henrissat B. The abundance and variety of carbohydrate-active enzymes in the human gut microbiota. Nat Rev Microbiol. 2013;11(7):497–504.

13. Danchin A. Bacteria in the ageing gut: did the taming of fire promote a long human lifespan? Environ Microbiol. 2018. [Epub ahead of print].

14. He Y, Wu W, Zheng HM, Li P, McDonald D, Sheng HF, et al. Regional variation limits applications of healthy gut microbiome reference ranges and disease models. Nat Med. 2018.

15. Deschasaux M, Bouter KE, Prodan A, Levin E, Groen AK, Herrema H, et al. Depicting the composition of gut microbiota in a population with varied ethnic origins but shared geography. Nat Med. 2018.

16. Ayeni FA, Biagi E, Rampelli S, Fiori J, Soverini M, Audu HJ, et al. Infant and Adult Gut Microbiome and Metabolome in Rural Bassa and Urban Settlers from Nigeria. Cell Rep. 2018;23(10):3056–67.

17. Lindeberg S, Jonsson T, Granfeldt Y, Borgstrand E, Soffman J, Sjostrom K, et al. A Palaeolithic diet improves glucose tolerance more than a Mediterranean-like diet in individuals with ischaemic heart disease. Diabetologia. 2007;50(9):1795–807.

18. Jonsson T, Granfeldt Y, Ahren B, Branell UC, Palsson G, Hansson A, et al. Beneficial effects of a Paleolithic diet on cardiovascular risk factors in type 2 diabetes: a randomized crossover pilot study. Cardiovasc Diabetol. 2009;8:35.

19. Whalen KA, McCullough ML, Flanders WD, Hartman TJ, Judd S, Bostick RM. Paleolithic and Mediterranean Diet Pattern Scores Are Inversely Associated with Biomarkers of Inflammation and Oxidative Balance in Adults. J Nutr. 2016;146(6):1217–26.

20. Otten J, Stomby A, Waling M, Isaksson A, Soderstrom I, Ryberg M, et al. A heterogeneous response of liver and skeletal muscle fat to the combination of a Paleolithic diet and exercise in obese individuals with type 2 diabetes: a randomised controlled trial. Diabetologia. 2018;61(7):1548–59.

21. De Filippis F, Pellegrini N, Vannini L, Jeffery IB, La Storia A, Laghi L, et al. High-level adherence to a Mediterranean diet beneficially impacts the gut microbiota and associated metabolome. Gut. 2016;65(11):1812–21.

22. Girard C, Tromas N, Amyot M, Shapiro BJ. Gut Microbiome of the Canadian Arctic Inuit. mSphere. 2017;2(1).

23. Dall’Asta C, Scarlato AP, Galaverna G, Brighenti F, Pellegrini N. Dietary exposure to fumonisins and evaluation of nutrient intake in a group of adult celiac patients on a gluten-free diet. Mol Nutr Food Res. 2012;56(4):632–40.

24. Biagi E, Franceschi C, Rampelli S, Severgnini M, Ostan R, Turroni S, et al. Gut Microbiota and Extreme Longevity. Curr Biol. 2016;26(11):1480–5.

25. Klindworth A, Pruesse E, Schweer T, Peplies J, Quast C, Horn M, et al. Evaluation of general 16S ribosomal RNA gene PCR primers for classical and next-generation sequencing-based diversity studies. Nucleic Acids Res. 2013;41(1):e1.

26. Masella AP, Bartram AK, Truszkowski JM, Brown DG, Neufeld JD. PANDAseq: paired-end assembler for illumina sequences. BMC Bioinformatics. 2012;13:31.

27. Caporaso JG, Kuczynski J, Stombaugh J, Bittinger K, Bushman FD, Costello EK, et al. QIIME allows analysis of high-throughput community sequencing data. Nat Methods. 2010;7(5):335–6.

28. Edgar RC. Search and clustering orders of magnitude faster than BLAST. Bioinformatics. 2010;26(19):2460–1.

29. Haas BJ, Gevers D, Earl AM, Feldgarden M, Ward DV, Giannoukos G, et al. Chimeric 16S rRNA sequence formation and detection in Sanger and 454-pyrosequenced PCR amplicons. Genome Res. 2011;21(3):494–504.

