## Supplementary figures and images for "Gut microbiome response to a modern Paleolithic diet in a Western lifestyle context"

### Supplementary file 1

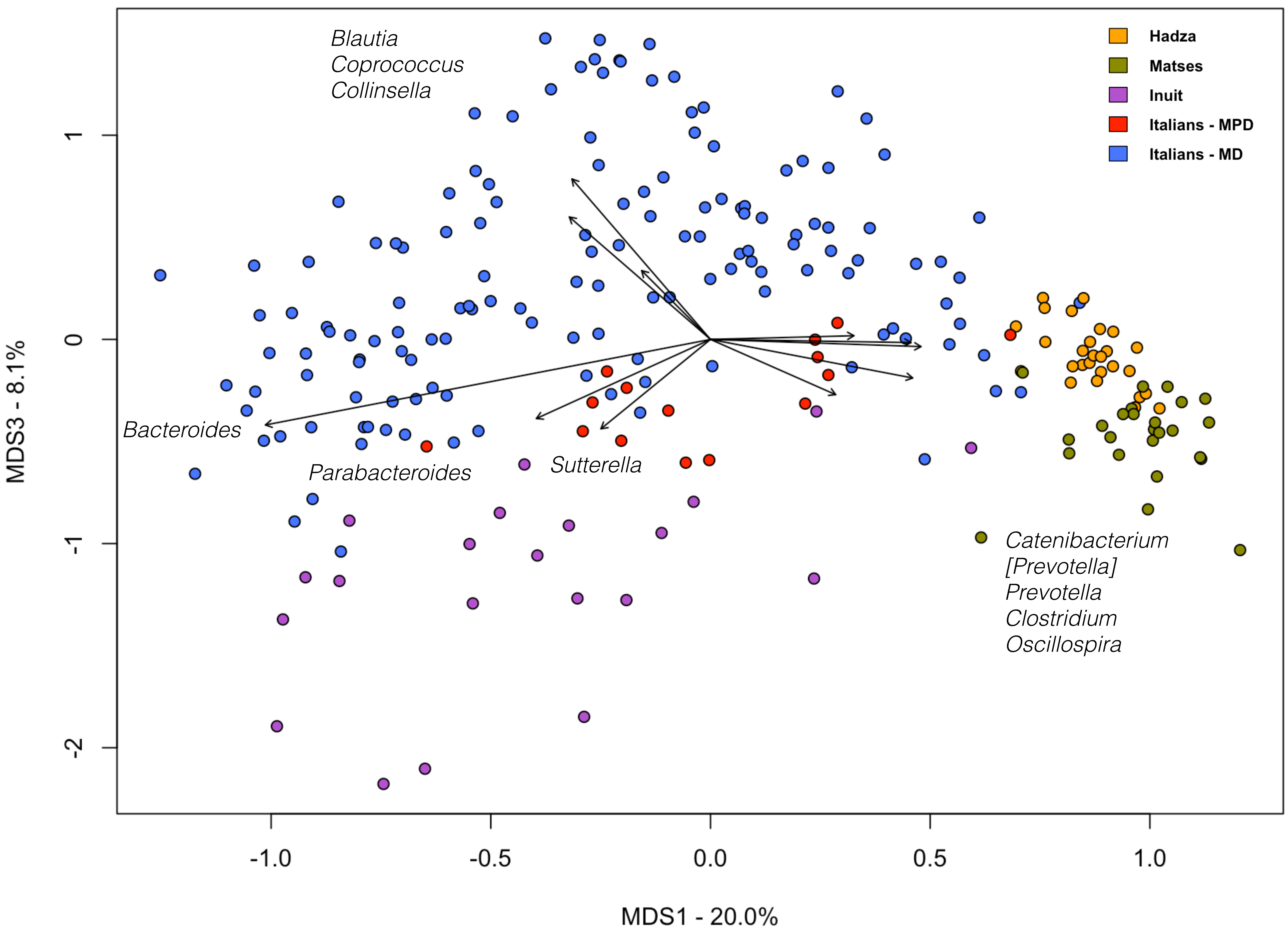
